# Inward thickening of embryonic neuron-packing brain walls: Mechanical integrity via curvature-associated inner-surface contractility

**DOI:** 10.64898/2026.05.01.722335

**Authors:** Arya Kirone Patra, Koyuki Inoue, Tomoki Nishikawa, Taisei Hiratsuka, Koichiro Tsujikawa, Kanako Saito, Takaki Miyata, Tomoyasu Shinoda

## Abstract

Despite recent explorations of tissue-level epithelial morphogenesis, how the embryonic brain wall, an epithelial derivative unique in its extensive cellular stratification coupled with epithelial-cell heightening to ∼0.5 mm, achieves thickening is unknown. Furthermore, the role of the inner curvature of the wall in this process remains unclear. Thus, whether the apically concave dorsal cerebral (pallial) wall thickens inward, not only outward as previously thought, was examined in culture and *in vivo*. The pallial wall in the midembryonic period, but not the earlier pallial wall, thickened inward at a pace equivalent to that of neuronal accumulation, which was found via stress-release tests to proceed in a compressive manner. The inhibition of actomyosin-mediated contraction of the inner/apical surface prevented the pallium from thickening, while inducing apical-surface buckling, suggesting the necessity of this inward thickening to actively avoid overcrowding. In contrast, inward bulging of the apically convex ganglionic eminence occurs more passively via pushing of the low-actomyosin apical surface by the constituent cells.

## Introduction

In tubes or boards that are physically bent for manufacturing and construction purposes, the material that runs the inner concave arc (intrados) of the bent tube is compressed and requires care so as not to cause deformations such as wrinkles, kinks, and buckles [1, 2]. Purposeful thickening of the wall of an already bent tube might be technically difficult or may not be the aim in manufacturing or fabrication processes. However, such thickening is an important morphogenetic task during brain development.

Embryonic epithelial tubes, formed by the lateral assembly of apicobasally polarized cells, exhibit inner/apical-surface concavity and undergo key morphogenetic growth processes, such as extension, branching, and ballooning [3–5]. Such dynamic transformations mostly occur while an epithelium maintains its original wall thickness (∼20 μm) or thickens only modestly (∼100 μm) through pseudostratification (i.e., cell cycle-dependent staggered nuclear positioning by epithelial cells) [6]. In contrast, developing brain walls are unique in that they undergo extensive thickening (∼500 μm; in mice) with the elongation of epithelial-like (apical-touching) neural progenitor cells (NPCs; also called radial glial cells; ∼500 μm spanning the wall), accompanied by an increase in the number of neurons that are generated in the inner zone of the wall to subsequently acquire nonepithelial (apical-untouched) characteristics [7]. Because neurons migrate outward and accumulate in the outer zones (Fig. 1A, red) [8–10], it is generally thought that dome-shaped (apically concave) walls such as the pallium (dorsal cerebrum) thicken outward. However, recent studies have shown that the lateral ventricle encompassed by the pallial wall narrows, with advancing development, toward the perinatal period in mammalian embryos [11–13], prompting us to investigate the possible inward thickening of the pallial wall and its biological significance or necessity.

**Figure 1.**
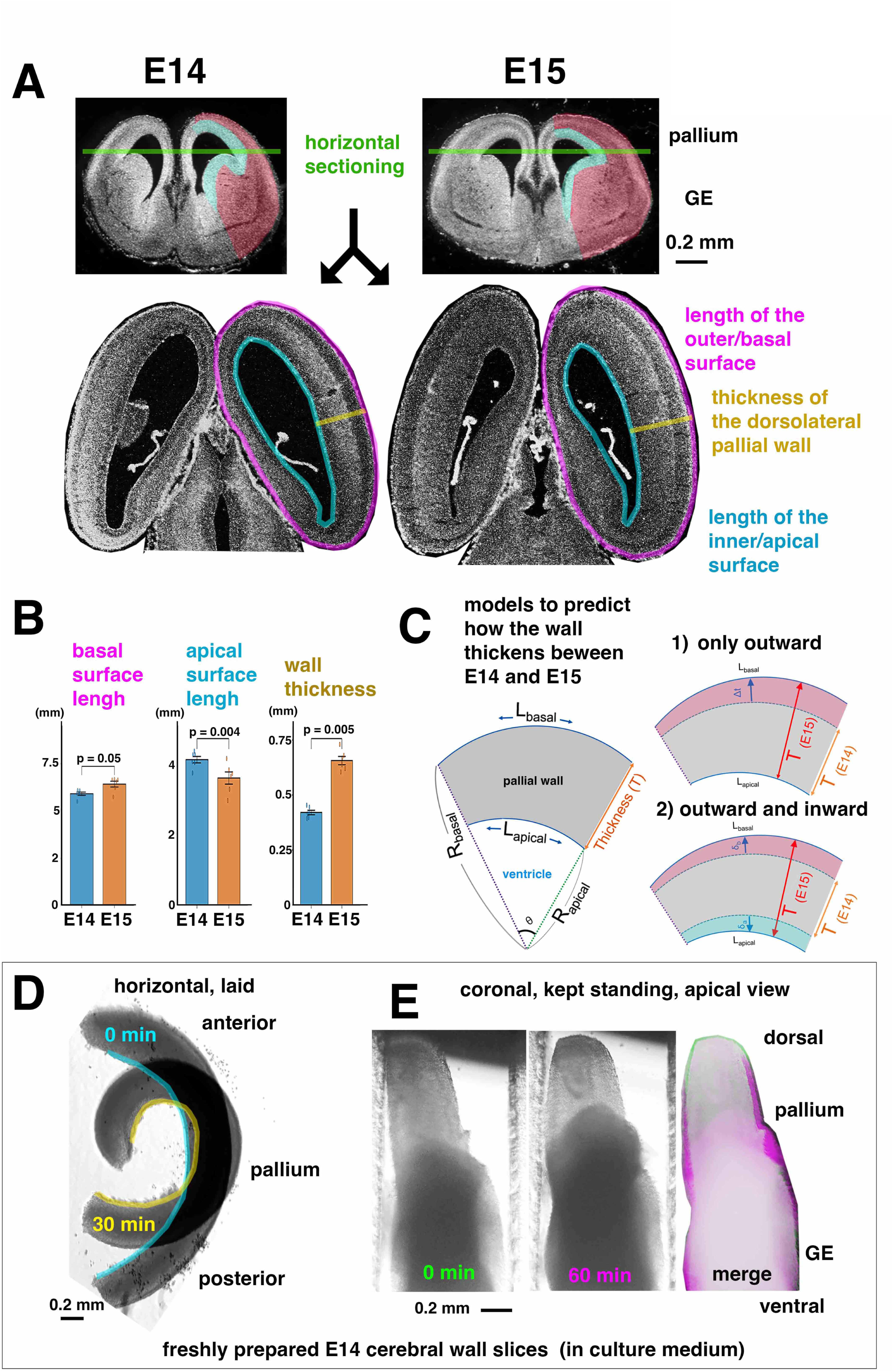
Physical factors that may be involved in the inward thickening of the E14 mouse cerebral wall. (A) Sections of the mouse cerebral wall were obtained between E14 and E15, and thickening of the wall was assessed. GE, ganglionic eminence. (B) Graph comparing the length of the outer/basal surface (Brunner‒Munzel test), the length of the inner/apical surface (Brunner‒Munzel test), and the thickness of the wall (Mann‒Whitney U test) in horizontal sections between E14 and E15. n=6 animals. Error bars, SEMs. Effect size (Cliff’s delta): 0.72, 0.77, and 0.69 respectively (large effect). (C) Models for measurement-based prediction of the pattern of wall thickening (see Methods for details). (D) A horizontally sliced E14 pallial wall that bent/curled inward during live observation. (E) A coronally sliced E14 cerebral wall (from the pallium to the GE) that remained upright with its inner/apical top surface. Note the lateral/circumferential spreading (magenta) of the wall’s constituent cells (nonepithelial, neurons) by 60 min.

Our analysis of the mouse pallial wall, whose inner curvature is concave, revealed that pallial wall thickening occurs not only outwardly but also inwardly in a stage-dependent manner, with frequent inward thickening at E14 and later. By combining surgical release of the prestress that exists *in vivo*, the observation of wall thickening dynamics in slice culture, and the inhibition of actomyosin-dependent contractility/narrowing of the wall’s apical/inner surface, we found that bidirectional thickening—both outward and inward—is necessary for the apically concave pallial wall to secure its mechanical integrity during the midembryonic period. This process ensures that constituent cells accumulate safely even under externally provided confining/compressive forces, such as one from brain-surrounding connective tissues [14, 15].

Furthermore, we report that the prospective striatal wall, which ventrally neighbors the pallium and has become apically convex by midembryonic days (thus called the ganglionic eminence [GE]), undergoes a different style of apical-surface management. Compared with the inner surface of the pallial wall, the inner surface of the GE wall shows weaker expression of F-actin and phosphorylated myosin light chain (pMLC) and is less contractile. As a result, the inner surface of the GE is more permissive to the wall’s constituent cells that push the inner surface, allowing the GE to further passively bulge into the ventricular space.

## Results and Discussion

### *In vivo* comparison at E14 and E15 supports a model of the inward thickening of pallial walls

As a first step to determine whether the midembryonic pallial wall, whose inner/apical surface is concave, thickens inward and, if so, how it achieves such inward thickening, we compared horizontal sections of the midembryonic pallial wall between E14 and E15 (Fig. 1A). As the wall thickened (from 0.420 ± 0.010 mm [n=6] to 0.656 ± 0.018 mm [n=6]), the total length of the outer/basal surface (magenta) increased (from 5.879 ± 0.088 mm to 6.385 ± 0.152 mm; p = 0.0478), and that of the inner/apical surface (cyan) decreased (from 4.151 ± 0.091 mm to 3.627 ± 0.173 mm; p = 0.00373) (Fig. 1B). We then used these measurements to predict the contribution of the inward shift—which can be inferred from the narrowing of the inner/apical surface—to the overall thickening of the wall (Fig. 1C). These results support a model in which the wall thickens bidirectionally (i.e., both outwardly and inwardly) (model 2) rather than one in which thickening occurs only outwardly (model 1) (see Supplemental Information).

Although it is possible that the predicted narrowing of the apical surface between E14 and E15 is due to the detachment of NPCs from the apical surface [16], a previous count suggested that the total number of apical endfeet (ZO1-immunostained NPCs) increased from E14 (1,344,000 per hemisphere) to E15 (1,768,000) [17], with an earlier decrease from E12 (1,892,000) to E13 (1,795,000) and to E14 and further increase from E15 to E16 (2,448,000). Therefore, NPC detachment alone cannot explain the narrowing of the apical surface from E14 to E15.

### Two mechanical factors for consideration in possible inward wall thickening

To further investigate the possible involvement of other factors, especially mechanical factors, we first examined horizontally prepared E14 pallial wall slices to determine whether they spontaneously bend/curl inward, as coronally prepared E14 pallial walls do so in an actomyosin-dependent manner [18, 19]. After 30 min, the contractility of the apical surface was clear, with narrowing/shortening to 75–80% of the original length observed (Fig. 1D).

Next, we used a surgical stress-release test to explore the *in vivo* mechanical conditions that had been formed by the nonepithelial constituent cells of the wall (i.e., mainly neurons). A recent pilot study of surgically prepared E14 pallial wall cubes (≈ 0.3 mm × 0.3 mm × 0.3 mm) revealed that neurons, which are nonepithelial (apical-unconnected) in nature, bulged out laterally (i.e., recoiled) along the cut edges in 30 min, increasing the width of the cube while decreasing its height [20]. To assess the likely pre-existing compressive packing of neurons at a more expanded spatial scale, we monitored coronally prepared E14 cerebral wall slices (0.5 mm thick; continuous from the pallium to the GE), focusing on their edges (Fig. 1E). At 60 min, lateral/circumferential recoil of neuron-like constituent cells was recognized at both the anterior and posterior edges of the coronal slice (magenta in the merged images), suggesting that the *in vivo* neurons had been circumferentially compressed until their surgical release and recoil.

### Cultured midembryonic pallial walls, but not pallial walls at earlier timepoints, thickened inwardly in coordination with neuronal accumulation

To determine whether freshly prepared E14 pallial walls thicken inward and, if so, how the contractile apical surface and the constituent neurons participate in this process, coronally prepared slices embedded in collagen gel were subjected to bright field and fluorescence imaging (Fig. 2; Movie 1A-C). Not only “open-type” coronally prepared slices (C-shaped; circumferential continuity lost) but also “closed-type” slices (ring-shaped; circumferentially continuous and geometrically closer to *in vivo* conditions than the “open-type” slices were) showed inward thickening at 12 h. A trend was also observed: slices with a greater initial thickness tended to show greater inward thickening (Fig. 2A). This tendency was more clearly observed when we compared midembryonic pallial walls with pallial walls at earlier timepoints (Fig. 2B; Movie 1A). Slices prepared at E14 (originally 0.3 mm or thicker) thickened inward (+ 11.6 ± 2.9% from the original thickness [n=4]) in 6 h, while their outward thickening was minimal (+ 1.8 ± 1.4%). E15 slices showed similar apically dominant thickening (+ 10.4 ± 7.8% apically and 0% basally [n=4]). Thickening of the slices prepared at E11 (originally 0.15–0.2 mm apicobasally) (n=5) was minimal both apically (+ 2.5 ± 1.9%) and basally (+ 3.4 ± 3.9%). The apicobasal thickness of the E12 slices (n=3) increased slightly (+ 5.3 ± 0.3% apically and + 5.4 ± 6.8% basally). E13 slices (n=4) showed initial signs of apically dominant thickening (+ 10.5 ± 9.1% apically and + 4.1 ± 4.9% basally).

**Figure 2.**
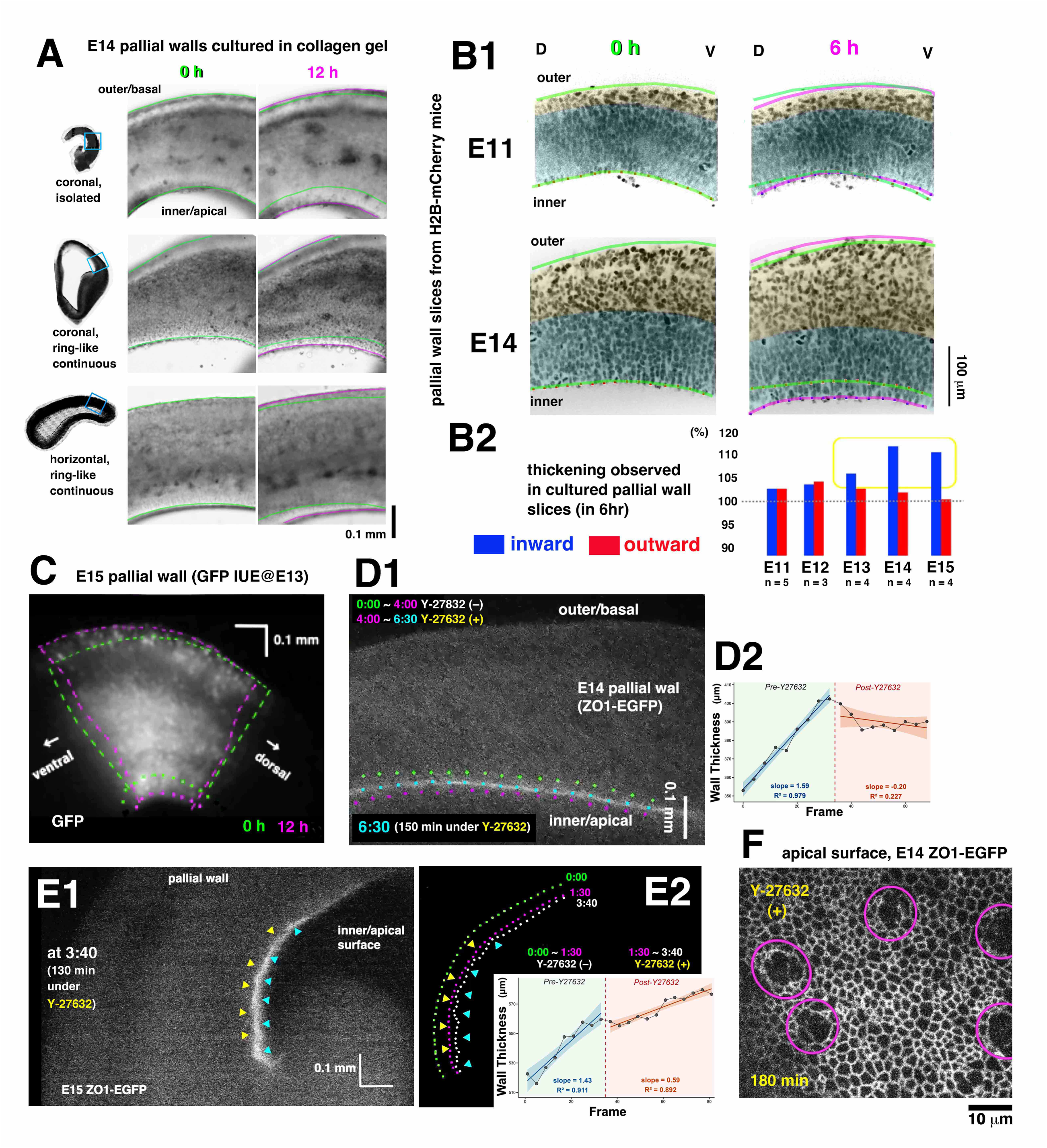
Cultured pallial walls thickened inward in coordination with neuronal accumulation under actomyosin-mediated contraction of the apical surface. (A) Bright-field time-lapse images of differently prepared E14 pallial wall slices (top, coronally sliced with the loss of circumferential continuity; middle, coronally sliced with ring-like continuity; bottom, horizontally sliced with maintained continuity). (B1) Fluorescent live-field images of pallial walls (coronally sliced from E11 or E14 H2B-mCherry mice in which nuclei can be visualized). (B2) Graph comparing the degree to which cultured pallial walls thickened toward the outer/basal or inner/apical sides. (C) Fluorescent live-field images of the E15 pallial wall labeled with GFP *in utero* at E13. (D1) Fluorescent live-images of the pallial wall (coronal, E14 ZO1-EGFP mouse for apical surface visualization) after the application of Y-27632 (20 μM) for 4 h. Five-minute interval. (D2) Wall thickness before and after Y-27632 treatment over time. (E1) Fluorescent live-field images of the pallial wall (coronal, E15 ZO1-EGFP) after the application of Y-27632 (20 μM) for 1.5 h. (E2) Trace of the apical surface before and after Y-27632 treatment to compare the position and curvature of the apical surface, coupled with a time series graph of wall thickness. Five-minute interval. (F) *En face* (apical-top) live-field images of the E14 ZO1-EGFP pallial wall that formed abnormally large holes (magenta circles) in the presence of Y-27632.

Remarkably, the inward thickening of the cultured E14–E15 pallial wall (i.e., an inward shift of the apical surface) was accompanied by thickening of the neuronal zone. In a representative E14 slice prepared from an H2B-mCherry mouse [21], the neuronal zone could be recognized as an outer, nonpseudostratified (nonepithelial-like) zone (Fig. 2B; Movie 1A). The number of H2B-mCherry-labeled nuclei increased from 331 to 434 (+31%) in the neuronal zone (from 92 to 119 in the most superficial “cortical plate” formed by postmigratory neurons; from 239 to 315 in the “intermediate zone” filled with migrating neurons). The number of nuclei in the pseudostratified zone (called the “ventricular zone”) marginally decreased from 355 to 321 (–11%). When E15 pallial walls that had been electroporated locally with GFP-expressing vectors *in utero* at E13 were cultured, a GFP-labeled fan-shaped (apically narrow, basally wide) region increased in apicobasal thickness and reduced in circumferential width by 12 h (n=5/5) (Fig. 2C; Movie 1C). Because the GFP-labeled fan-like region was sandwiched between more dorsal and ventral regions (Movie 1C, bright field), its dorsoventral narrowing may be associated with a circumferentially compressed status, inferred from progressively accumulating neurons (Fig. 1E). Notably, the dorsoventral narrowing of the neuron-rich fan-shaped labeled region was spatiotemporally coordinated with (1) narrowing of the apical surface and (2) wall thickening in both directions.

### The inhibition of actomyosin prevented the apical surface from shifting inwardly, causing irregularities such as wrinkles and holes

To determine whether and, if so, how tangential contractility of the apical surface contributes to the inward thickening of midembryonic pallial walls, pharmacological experiments were performed using Y-27632, a rho kinase inhibitor that is known to inhibit actomyosin-dependent apical surface contractility [18, 22, 23]. Our preparative examinations revealed that the apical surface, which should have narrowed (detected as shortening) to 75–80% of the original length in free isolated slices in 30 min (Fig. 1D), narrowed to only 90% under Y-27632 treatment. When E14 pallial wall slices prepared from ZO1-EGFP mice [24] cultured in collagen gel were exposed to Y-27632, the inward shift of the apical surface (fluorescently labeled) that was observed during the pre-Y-27632 period quickly and completely ceased within 5 min in a highly reproducible manner (n=18/18), followed by slight retraction (from ∼15 μm/h apically to ∼4 μm/h basally) of the apical surface (Fig. 2D; Movie 2A). Furthermore, E15 slices treated with Y-27632 similarly stopped shifting or exhibited severe deceleration (from ∼15 μm/h to ∼6 μm/h apically) of the inward shift of the apical surface (Fig. 2E; Movie 2B).

In addition to the deceleration or disappearance of the inward shift, the Y-27632-treated apical surface displayed irregularities in curvature and endfeet meshwork. Fig. 2E (and Movie 2B) shows the emergence of wrinkles in an E15 ZO1-EGFP slice. Similar or mildly wrinkled (zigzag-like pattern) apical surfaces were frequently observed in E14 and E15 ZO1-EGFP slices in a region-specific manner (n=3/3 at E14, 3/4 at E15 if the most ventral part of the pallial wall that is neighbored by the striatum was inspected). Previous studies have reported that subapical overcrowding is most easily inducible (via NPC overproliferation) in the pallium near the striatum [25, 26], probably because the circumferential spread of the pallial element seems to be physically blocked by the striatum, as previously suggested by Clark [27]. This spatial overlap between the zigzag-like phenotype caused by reducing the contractility of the apical surface (and thus less inward shiftability) and the previously reported susceptibility to overcrowding suggests that the apical surface may normally play a role in spatial management of the wall’s constituent cells, most directly for the inward shift of subapical cells but also indirectly for the positioning of other cells along the apicobasal width. The wrinkled or zigzag-like irregularity shown by the less contractile/constrictable apical surface suggests physical buckling of the apical surface [28, 29], akin to the formation of villi through a relative abundance of the epithelial layer that occurs tangentially owing to narrowing of the adjacent mesenchyme [30]. *En face* live-field imaging further revealed that when Y-27632 was added, the ZO1-EGFP-labeled apical-surface meshwork of E14 pallial walls retracted basally at 2∼5 μm/h, recognized by progressive image blurring that required manual refocusing, and some apices became abnormally large (∼10x) (Fig. 2F; Movie 2C).

### *In vivo* Y-27632 experiments reproduced apical-surface buckling and thus supported the inward shift model

To determine whether the apical-surface phenotypes observed in pallial wall slices exposed to Y-27632 in culture could be reproduced *in vivo*, we injected Y-27632 into the lateral ventricle of E14 –E15 mice (at a final concentration of 100 μM) (Fig. 3A). Three hours after injection, clear wrinkling of the apical surface was detected in 72.2% of the Y-27632-treated mice (n=18) (Fig. 3B–E), whereas 13.3% of the control mice (DMSO-injected; n=15) showed minor apical-surface irregularity (p = 0.00128; Fisher’s exact test) (Fig. 3E). Possible fixation-induced tissue shrinkage may explain the minor irregularity observed in a small population of control samples. Wrinkling was preferentially found in the ventral-most pallium neighboring the GE, as in the slice culture, and in the dorsal-most pallium, where the curvature was greatest. Notably, the basal border of the pseudostratified zone (consisting mainly of NPC nuclei) also showed irregularities (zigzag-like patterns) with basal bumping at the dorsoventral level where the apical surface exhibited sharp, valley-like invaginations (Fig. 3B, 3C), suggesting buckling of the subapical/pseudostratified zone coupled with that of the apical surface. These findings therefore strongly suggest that tangential narrowing of the apical surface that should automatically shift it inward when its curvature is concave was insufficient, providing us with solid *in vivo* evidence that the midembryonic pallial wall would otherwise (i.e., normally) indeed thicken inward.

**Figure 3.**
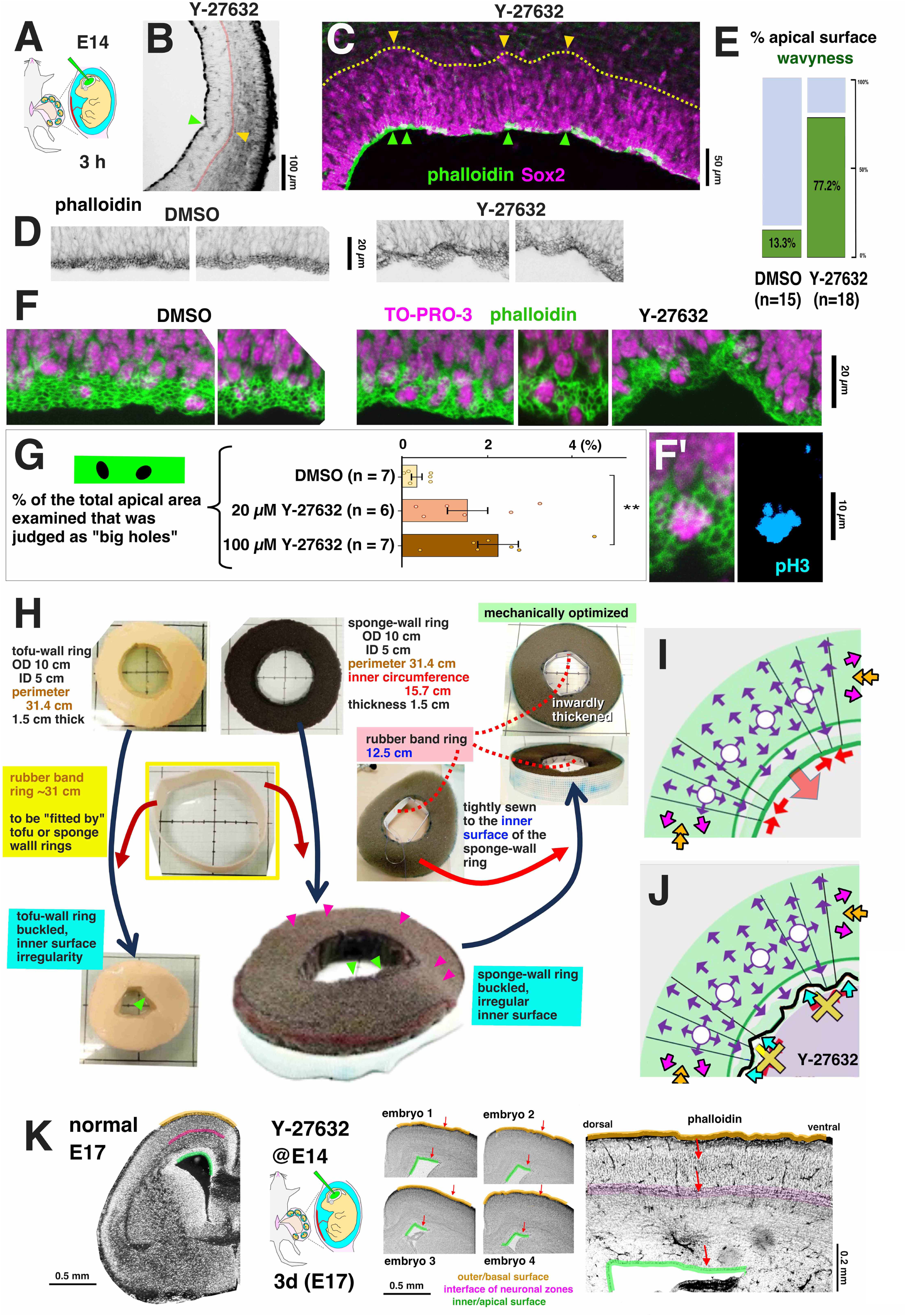
(A) Actomyosin-dependent inward thickening as a mechanical solution for overcrowding due to neuronal accumulation in the midembryonic pallial wall. Y-27632 was intravenously injected into E14 embryos *in utero*, followed by fixation at 3 h after injection. (B, C) Phalloidin photomicrographs showing the abnormally wrinkled inner/apical surface of the Y-27632-treated pallial walls. Green arrow, invagination of the apical surface. Yellow arrow, bulging of the basal border of the pseudostratified zone (filled with Sox2^+^ nuclei, C). (D) Magnified photomicrographs showing the wavy apical surface of Y-27632-treated embryos. (E) A mosaic graph comparing the frequency with which the wrinkled pattern was observed assessed using Fisher’s exact test. p = 0.00128. Odds ratio = 0.07. CI [0.005, 0.441]. (F) Costaining with phalloidin (green) and TO-PRO-3 (magenta). (F’) Costaining with phalloidin (green) and anti-pH3 (cyan). (G) Graph comparing the frequency with which large holes were observed. Kruskal‒Wallis rank-sum test followed by the Steel–Dwass post hoc test (DMSO vs. Y-27632 (final concentration of 20 μM) vs. Y-27632 (final concentration of 100 μM)). n = 6–7 animals. Error bars, SEMs. (H) Physical simulation using tofu and sponge walls and elastic rubber bands (see text for details). (I, J) Schematic diagrams comparing normal and Y-27632-treated pallial walls. (K) Bright-field photomicrograph of a coronally sectioned normal E17 cerebrum (left) and phalloidin photomicrographs of E17 mice that received Y-27632 via injection at E14 (right).

Abnormality on the apical-surface meshwork was also reproduced *in vivo* (Fig. 3F, G). We found that large holes (∼100 μm^2^) formed at M-phase NPC cell bodies (Fig. 3F’). This finding is quite similar to observations in chicken embryos treated with Y-27632 during neural tube closure [23]. In normal pallial walls at E14, M-phase cell bodies, even maximally rounding up, are deeper than the apical surface [19], and the average apex size of M-phase NPCs is 12.1 μm^2^, which is only slightly larger than the average of all apices (8.3 μm^2^) [17]. These abnormally large holes (∼100 μm^2^) suggest that the normal spatial relationship, as well as mechanical balance, between M-phase cell somata and the apical surface was disrupted. Healthy centripetal narrowing of the apical surface coupled with apical-ward pulling of the subapical structure may “counteract M-phase apical expansion [23]”, physically preventing M-phase cells’ somata from being exposed to the ventricular lumen.

### Mechanical simulation suggests that inward wall thickening is necessary for mechanical integrity

We then performed a physical simulation using a ring-shaped tofu wall or sponge wall (flat, 1.5 cm thickness, 10 cm outer diameter) to represent the pallial wall (Fig. 3H). Mathematical vertex models have suggested that apical constriction plays a role in maintaining the smoothness of the concave curvature upon the proliferation of epithelial cells [31]. We sought to address the apparent contribution of the apical surface to wall thickening, which may proceed under spatial limitations. During the midembryonic period, connective tissues surrounding the brain undergo differentiation into ossification [14, 15], making these tissues stiffer and less elastic, as demonstrated by a microsurgical tissue-recoil test [32]. When this mild centripetal/inward constriction was mimicked using an elastic rubber band, into which a ring-shaped tofu wall as a model of pallial walls was fitted at a certain degree of tightness, the inner surface of the tofu ring became wavy. Similarly, inward-confined sponge wall rings also could not maintain the original completely flat geometry, resulting in uneven or wavy surfaces, which is indicative of an overabundance of tissue volume than can be safely accommodated. *In vivo*, an increase in wall volume (the number of constituent cells) via cell production would also generate an analogous problem unless this increase was managed. Remarkably, this “overabundance” phenotype was rescued (i.e., the sponge wall became flat again) when another elastic/contractile rubber band ring (smaller than the inner circumference of the sponge-wall ring) was securely sewn onto the inner surface of the sponge-wall ring to tangentially narrow it, thereby pulling it centrally.

These simulation results therefore suggest that the active (energy-consuming) inward shift of the apical surface in the midembryonic pallial wall plays an essential role in safely managing the wall’s tissue volume and cytoarchitecture, preventing overcrowding (Fig. 3I). If this wall-thickening mechanism under mechanical integrity based on apical-surface contractility is inhibited, primary malformations along/near the inner surface such as wrinkles or buckles, quickly emerge (Fig. 3J). Once buckling has occurred, losing normal continuous concavity, the resulting mechanical instability, as revealed in manufacturing studies [1, 2], seemed to have produced the secondary and/or cumulatively-formed abnormalities involving the entire wall components by E17 (n=4 of 4 embryos that had been treated with Y-27632 at E14). In these E17 mice, the pallial wall’s dome-like structure was compromised, exhibiting abnormally straightened or bulged apical surface in the dorsolateral part together with sharp-angled bending of the dorsal-most part. Its basal surface, as well as the underlying neuronal layers, showed indentation(s) or depression(s) in the dorsolateral part (Fig. 3K). Thus, making inward room for midembryonic (E14) pallial wall’s constituent cells is necessary for the subsequent progression of appropriate histogenesis along both the radial axis and the circumferential axis until late embryonic days (E17), thereby underlying the proper circuit formation.

### The GE thickens inward via pushing of the low-actomyosin convex apical surface by the constituent cells of the wall

In our time-lapse monitoring, the prospective striatal wall, whose inner/apical surface is convex (thus called the ganglionic eminence [GE] during the embryonic period), also thickened inward (Fig. 4A; Movies 1B, 3A, and 3B). However, the response of the apical surface to Y-27632 application differed from that of the pallium at E14. While retraction of the pallial apical surface upon Y-27632 treatment suggested an active, inward shift driven by narrowing of the wall, the GE apical surface continued to shift inwardly even after Y-27632 application (Fig. 4A), and bumps sometimes formed (Movie 3B).

**Figure 4.**
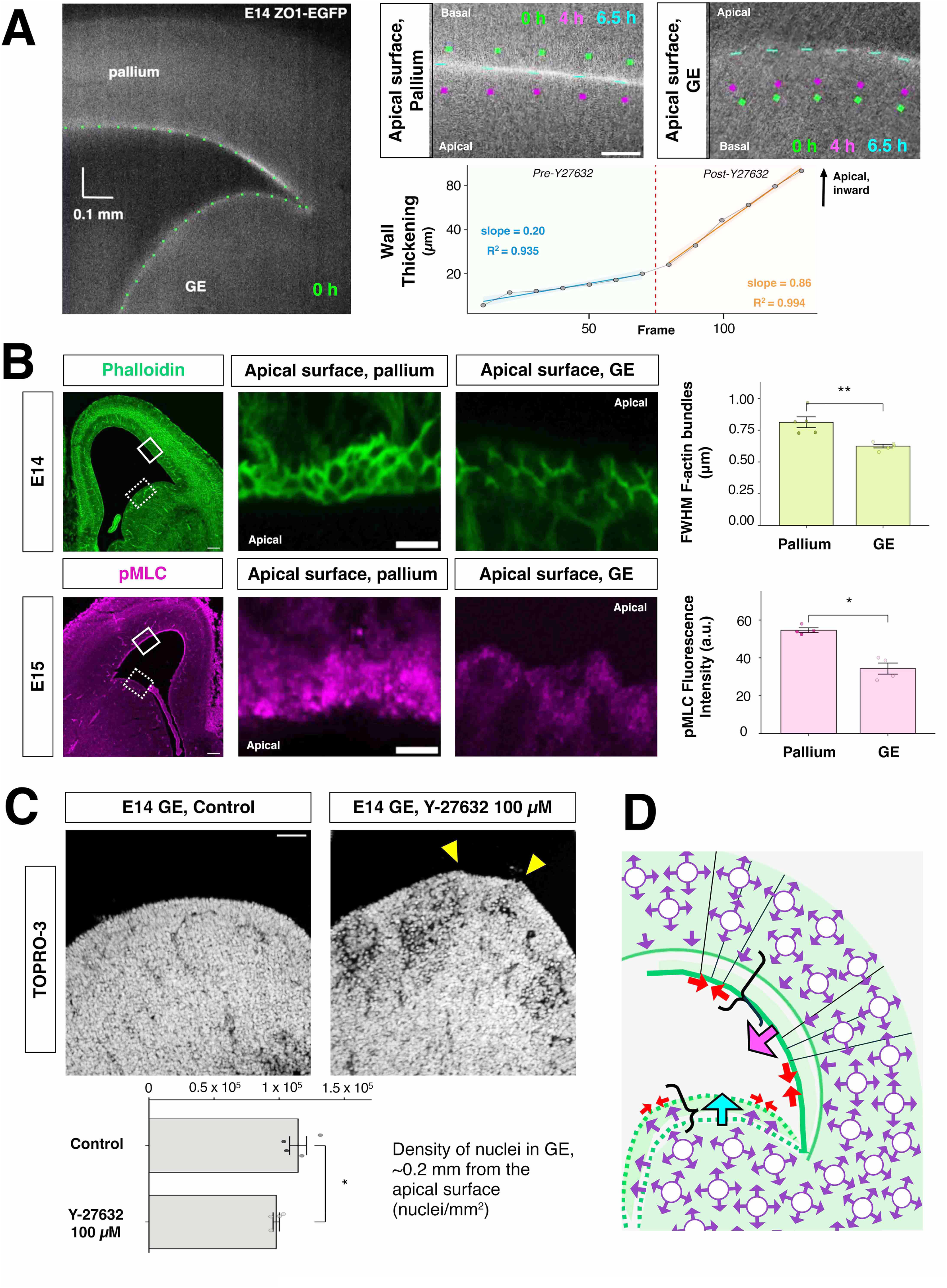
The midembryonic ganglionic eminence also grows inward in a manner that mechanistically differs from that of the pallial wall. (A) Fluorescent live-field images of cerebral (pallial and GE) walls (coronal, E14 ZO1-EGFP) after the application of Y-27632 (20 μM) for 4 h, accompanied by a time series graph showing the inward shift of the apical surface before and after Y-27632 treatment (baseline measurement from the position of the apical surface at Time = 0 min). 1 Frame = 5 min. Bar, 50 μm. (B) Comparison of phalloidin [green; E14_Pal_ = 0.812 ± 0.0429 vs E14_GE_ = 0.625 ± 0.0141 (mean ± SD)] and phospho myosin light chain (pMLC) [magenta; E15_Pal_ = 54.622 ± 1.268 vs E15_GE_ = 34.308 ± 2.916 (mean± SD)] staining between the pallial and GE walls. Bar, 100 μm for the left panels; 5 μm for the remaining panels. Mann‒Whitney U test. n = 5 and 4, respectively. Error bars, SEMs. Effect Size (Cliff’s delta) = 1 (both cases; large effect). \**p* < 0.05, ** *p* < 0.01. (C) Comparison of the morphology of the GE (top) and the density of the nuclei of cells in the GE (bottom graph) between normal and Y-27632-treated E14 embryos at 3 h. Bar, 50 μm. Mann‒Whitney U test. n = 4. Error bars, SEMs. Effect Size (Cliff’s delta) = 1 (large effect). * *p* < 0.05 (D) Schematic illustration of the differential contributions of the apical surface to the mechanical integrity underlying wall thickening, dependent on its contractility and wall geometry (curvature).

A previous study revealed that spontaneous narrowing of the apical surface in isolated slices (Fig. 1D) was more extensive in the pallium than in the GE [33]. That study also showed that tangential recoils of the apical surface upon laser ablation were smaller in the pallium than in the GE (in terms of displacement and velocity), but the degree of reduction in recoil by actomyosin inhibition was greater in the pallium than in the GE, suggesting that tension on the apical surface is generated differently: in the pallium, active (actomyosin-dependent) centripetal narrowing of an apex pulls the surrounding apices centrally (NPCs intrinsically narrow, generating tension), whereas the apical surface of the GE is more passively stretched, acquiring tension via apical-ward pushing from the constituent cells of the GE.

To determine whether this difference between the pallium and GE with respect to the mechanical conditions of the apical surface, as revealed by Y-27632 treatment and laser ablation tests, also explains their differential inward thickening, we compared F-actin visualized by phalloidin staining and immunohistochemically detected pMLC between the pallium and the GE (Fig. 4B). Our measurement of the apical-surface meshwork revealed that the expression of F-actin and pMLC per unit apex–apex border (corresponding to the cell‒cell junction) was greater on the pallial apical surface than on the GE apical surface (Fig. 4B), supporting the abovementioned model. Consistent with these findings, intraventricular injection of Y-27632 induced GEs to adopt slightly more bulged or bumpy shapes, where the density of GE-containing cells was lower (more dilute per unit space/area, 0.97×10^5^ ± 2394.2 cells/mm^2^) than that of control GEs (1.14×10^5^ ± 6449.5 cells/mm^2^, p=0.03038, Mann‒Whitney U test) (Fig. 4C). These results suggest that inward thickening of the GE wall can occur relatively passively compared with that of the pallium in a manner associated with the wall geometry (convexity) and weaker actomyosin of its apical surface.

How this dorsal-high-ventral-low actomyosin expression pattern is established in the developing mammalian brain remains unknown. As the ventral telencephalic tubes are still apically concave in the early embryonic period, actomyosin may be involved in this concave-to-convex transition; therefore, the factors that trigger such transitions and the mechanosensing mechanism that occurs in the prospective GE region should be investigated.

In summary, inward thickening of the pallial and GE walls can be abstracted as two different types of trains running down a gentle slope. With increasing numbers of constituent cells (neurons), analogous to passenger cars, the highly contractile pallial apical surface works as the leading locomotive with an engine brake—it gently resists the gravity-mediated free downward/forward acceleration provided by the load of the passenger cars and simultaneously actively drives the entire train forward at a pace it determines itself. In contrast, the mechanically less active GE (future striatal) apical surface behaves more like a relatively freely rolling carriage, advancing inward under the pushing force of the GE’s own constituent cells. The growth of these two neighboring regions while maintaining their distinct inner-wall curvatures, i.e., concave and convex, based on differential apical-surface actomyosin, may achieve the following: (1) promoting space efficiency so as not to become competitive by allowing only the GE to utilize the lateral-ventricular space more easily/freely, (2) providing mechanical protection (dorsally) from the confining forces generated by the skull-forming tissue elements; and (3) supporting the coformation of laminar/cortical and nuclear structures, establishing a spatiotemporal basis for their postnatal functional collaboration.

## Acknowledgement

We thank M. Masaoka, N. Noguchi, J. Yamada for excellent technical supports; Miyata lab’s people for discussion. This work was supported by JSPS Grant-in-Aid 16H02457, 21H02656, 21H00363, 23H04306, 24K02205 (T.M.), 17K10176, 20K07224 (K.S.), 17K08489, 20K07223, 23K06319 (T.S.).

## Supplemental Information

### Materials and Methods

#### Animals

Animal experiments were conducted according to the Japanese Act on Welfare and Management of Animals, Guidelines for Proper Conduct of Animal Experiments (published by Science Council of Japan), and the Fundamental Guidelines for Proper Conduct of Animal Experiments and Related Activities in Academic Research Institutions (published by the Ministry of Education, Culture, Sports, Science and Technology of Japan). All protocols for animal experiments were approved by the Animal Care and Use Committee of Nagoya University (No. 29006). R26-H2B-mCherry transgenic mice (accession No. CDB0239K) [21, 25] and R26-ZO1-EGFP transgenic mice (accession No. CDB0260K) [19, 21] were provided by Toshihiko Fujimori (NIBB, Japan). Timed-pregnant ICR mice were obtained from SLC (Hamamatsu, Japan). The day when the vaginal plug was detected was considered E0.

### Bright-field imaging of free isolated cerebral walls to observe their spontaneous inward bending or tissue recoiling from their cut surfaces

Coronal or horizontal slices 0.3–0.5 mm thick) were manually prepared from embryonic mouse cerebral hemispheres (pallial and striatal regions together). All surgical procedures were made in a polystyrene culture dish (Corning) containing DMEM/F12 using microknives. The bottom of the medium-containing dish had previously been coated (5 mm thick) with silicone rubber made by mixing KE-103 and CAT-103 (Shin-Etsu Chemical, Tokyo, Japan).

Bending/curling was recorded using an Olympus IX71 (4×) equipped with an on-stage culture chamber (Tokai Hit), temperature-controlled and filled with 5% CO_2_, and an Orca ER camera (Hamamatsu Photonics), as previously described [19, 25, 26]. Quantitative measurement was performed using ImageJ. Movies were captioned or assembled using After Effects (Adobe).

### Fluorescent imaging of gel-mounted cerebral walls

Cerebral walls similarly sliced were embedded in type-I collagen gel (Nitta Gelatin) prepared in a cell culture dish (Corning) or a glass-bottom dish (Iwaki) at a concentration of 0.9–1.1 mg/ml. They were cultured in DMEM/F12 supplemented with 5% horse serum and 5% fetal bovine serum. Time-lapse confocal microscopy was performed using an Olympus IX71 (4×) equipped with an Orca ER camera (Hamamatsu Photonics), a CV1000 spinning disc confocal system (Yokogawa, Japan), or a BX51W1 microscope (Olympus) equipped with a CSU-X1 spinning disc confocal unit (Yokogawa) in an on-stage culture chamber (Tokai Hit) filled with 55% N_2_, 40% O_2_, and 5% CO_2_., according to the previously described methods [19, 25, 26]. At a certain point, Y-27632 was added to obtain a concentration of 20 μM.

### *In utero* electroporation

E13 mice were electroporated with pCAG-EGFP as previously described [25, 26]. After DNA solution was injected into the lateral ventricle, the head of the embryo in the uterus was placed between the discs of a forceps-type electrode (disc electrodes of 1 mm; CUY560P1, NEPA GENE, Chiba, Japan), and electric pulses (50 V) were administered four times, resulting in gene transfection into the cerebral wall.

### *In vivo* Y-27632 experiment

Pregnant mother mice were anesthetized via the intraperitoneal administration of a mixture of 0.75 mg/kg medetomidine hydrochloride (Orion Pharma, ZENOAQ), 4 mg/kg midazolam (SANDZ), 5 mg/kg butorphanol tartrate (Meiji Seika), and 1.5 mg/kg atipamezole hydrochloride (Kyoritsu Seiyaku). After midline laparotomy of the mother mouse, the uterine horn was pulled out. Then, a fine glass-needle filled with Y-27632 (Wako) (0.2–0.4 mM) or DMSO (Sigma-Aldrich) (1%) was inserted into the midbrain vesicle of embryos *in utero*, immediately followed by an injection of 0.5–1 μl, which was considered to provide a final concentration at 20–100 μM for Y-27632 according to our previous measurement of the total ventricular volume [34].

### Physical simulations

Based on the usefulness of physical (non-mathematical) models for simulating morphogenetic events during development [35], a simulation of physical events involving the midembryonic pallial wall (Figure 3H) was prepared using mechanically appropriate materials (e.g., rubber band rings to mimic the contractile apical surface of the pallial wall and the brain-surrounding connective tissues; sponge or tofu blocks to model a cerebral wall).

### Immunofluorescence

Embryonic mice (E14, 15, 17) were first transcardially perfused with 4% paraformaldehyde (PFA) or periodate-lysine-paraformaldehyde (PLP) fixative (whose paraformaldehyde concentration 2%) containing 1 μM Taxol and 0.1% Triton X-100. (4 °C 1∼3 h). Following which the embryonic brains were further fixed for 1.5 h at 4°C with PLP or 4% PFA fixatives containing 1 μM Taxol and 0.1% Triton X-100. For immunostaining of phospho-myosin light chain (pMLC), 1% trichloroacetic acid in PLP was used. Frozen sections (16 μm thick for coronal; 18 μm for horizontal) were treated with anti–phospho-myosin light chain (ab2480, Abcam, 1:500), anti–βIII-tubulin (mouse, MMS-435P, COVANCE, 1:1000), eFlour 660-conjugated anti-human/mouse Sox2 monoclonal antibody (Cat# 50-9811-82, InVitrogen, 1:300), or anti–phospho-histone H3 (rat, ab10543; Abcam, 1:500). After washes, sections were treated with TOPRO-3 (T3605, Thermo-Fisher. 1:1000), Alexa Fluor 488–conjugated phalloidin (A22283, Thermo Fisher Scientific, 1/1000), or Alexa Fluor 488–, Alexa Fluor 546–, or Alexa Fluor 647–conjugated secondary antibodies (Molecular Probes, A-11029, A-11006, A-11034, A-11030, A-11035, A-11081, A-21236, A-21245, 1:200). Images were obtained on an FV1000 laser scanning confocal microscope (Olympus, Tokyo, Japan). Images were arranged using Photoshop (Adobe).

### Mathematical modeling of wall thickening (Fig 1C)

We ask whether the E14→E15 thickening of the pallial wall occurs by outward basal shift alone (Model 1) or by bidirectional shift of both surfaces (basal and apical; Model 2). We fit an effective circular-arc geometry to E14 measurements, then, under the assumption that the angular extent θ is preserved between stages, use each model to predict E15 surface lengths and compare with the manually measured values (MMV) (see table for observations).

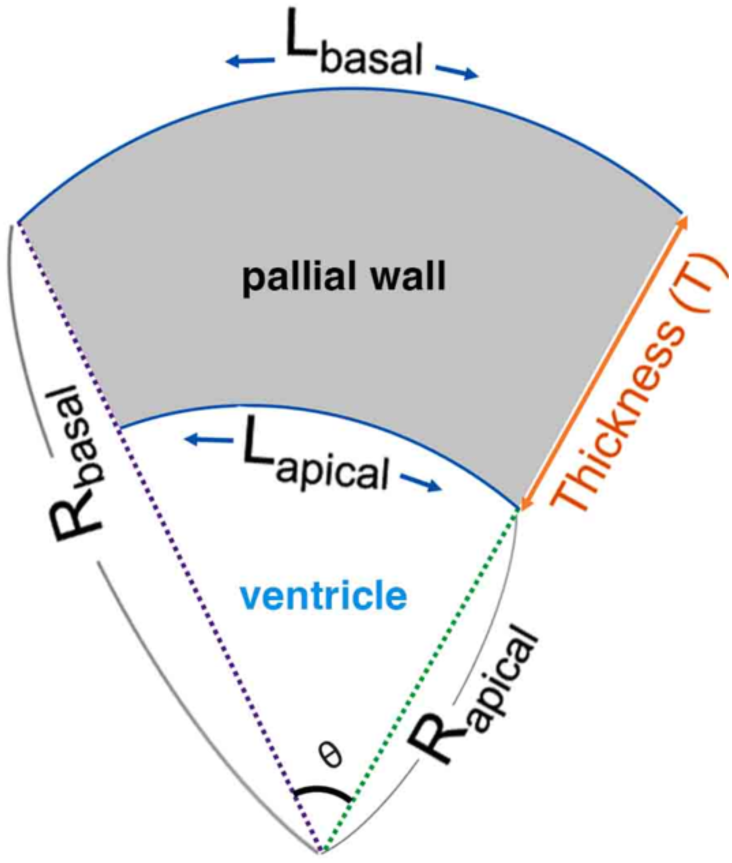

Definition of variables:

L_A_X_ = Length of the Apical surface at stage X

L_B_X_ = Length of the Basal surface at stage X

T_X_ = Thickness of the brain wall at stage X

R_A_X_ = Radius of Curvature of the Apical surface at stage X

R_B_X_ = Radius of Curvature of the Basal surface at stage X

ΔT= Difference in brain wall thickness between E15 and E14.

δ_A_ = Variable change (decrease) in R_A_ as thickness increases

δ_B_ = Variable change (increase) in R_B_ as thickness increases

Arc length (L) = Radius of Curvature (R) × Angle swept by Arc (θ)

Therefore,

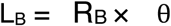

Since R_B_ = R_A_ + T, substituting values we get:

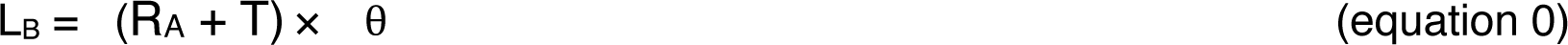

Now L_A_ = R_A_ × θ, putting value for R_A_ in (equation 0) and solving for θ we get:

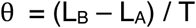

We used our MMV (see table; Fig. 1A, B) from E14 samples (L_A_E14_, L_B_E14_, and T_E14_) to define an effective geometry — essentially asking: “what circular arc would produce these exact apical and basal lengths with this exact thickness?”. The value of the angle swept (θ) came out to be 4.108 rad.

Table of Manually Measured Values (MMV)

**Table of Manually Measured Values (MMV)**
| Stage | Apical surface length (mm) | Basal surface length (mm) | Pallial wall thickness (mm) | Sample Size (n) |
| --- | --- | --- | --- | --- |
| **E14** | 4.151754 ± 0.091 | 5.879040 ± 0.088 | 0.420452 ± 0.010 | 6 |
| **E15** | 3.627290 ± 0.173 | 6.385941 ± 0.152 | 0.656590 ± 0.018 | 6 |

Further, we can get the values of R_A_E14_, from R_A_E14_ = L_A_E14_ /θ. We can also calculate R_B_E14_ as R_B_E14_ = R_A_E14_ + T_E14_. Hence, we get:

R_A_E14_ = 1.011 mm

R_B_E14_ = 1.431 mm

To verify the value of θ we check for the value of L_B_E14_ (L_B_E14_ = R_B_E14_ ×θ), which comes out to be very close to the actual value observed from the MMV.

Then we use these parameters to predict the values from E15. Here we ask, if the tissue thickens while preserving the same angular extent, what happens to the surface lengths?

#### Assumptions

1. The wall cross-section can be approximated as a circular arc.
2. θ is preserved between E14 and E15.
3. The wall is treated as having a single apical and a single basal radius of curvature.

In our view these are reasonable assumptions given the short time frame (24 hrs) and that the overall geometry of the brain wall does not change drastically in this period.

#### Model 1 [only Basal surface shifts]

This model assumes that the thickness is mediated by outward shift of the Basal surface alone. Therefore, the apical radius does not change and is equal to 1.011 mm at E15 (hence, R_A_E15_ = 1.011 mm). And since the angular extent also remains the same, θ = 4.108 rad.

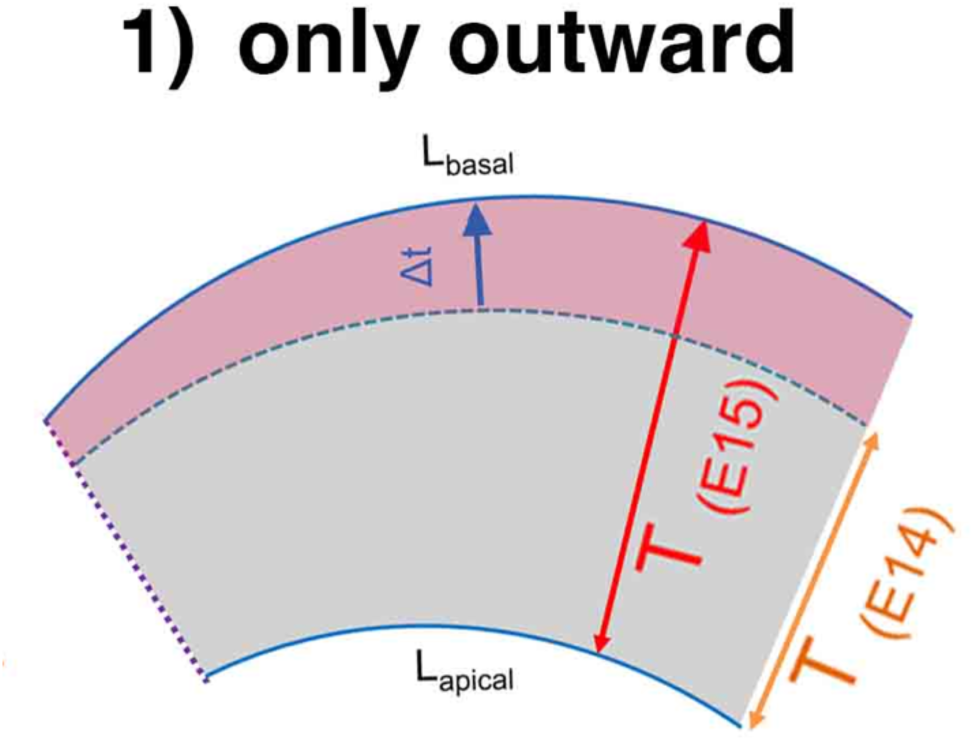

R_B_E15_ = R_A_E15_ + T_E15_ (we get T_E15_ = 0.656 mm, from MMV)

R_B_E15_ = 1.667 mm

Using this we calculate:

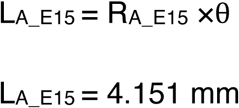

L_A_E15_ = 4.151 mm

And,

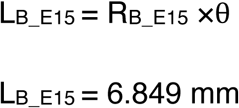

L_B_E15_ = 6.849 mm

Both values are larger than the observed values from MMV (see table). L_A_E15_ is overestimated by ∼14% and L_B_E15_ by ∼7%. The model fails badly on the apical side and moderately on the basal side, which rules out outward-only thickening.

#### Model 2 [Both Basal and Apical surface shift]

This model assumes that the thickness is mediated by a bidirectional shift—outward shift of the Basal surface and inward shift of the Apical surface. In this case, the apical radius and the basal radius change variably (δ_A_ and δ_B_) but the angular extent still remains the same, θ = 4.108 rad.

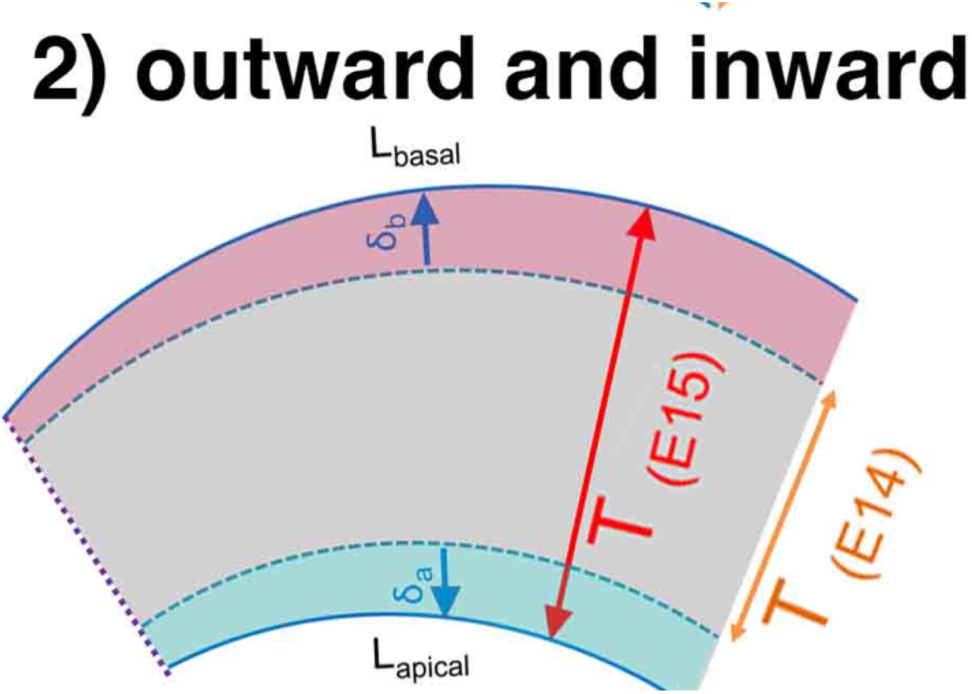

ΔT = T_E15 -_ T_E14_ = 0.236 mm (from MMV)

Since changes in radius are unknown:

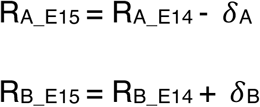

And,

δ_A_ + _B_ = ΔT (since the total change in surface radii must account for ΔT)

Using this we calculate:

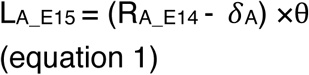

(Since the apical shift happens towards the center of the arc)

And,

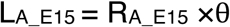

L_B_E15_ = (R_B_E14_ + δ_B_) ×θ(since the basal shift happens away from the center of the arc)

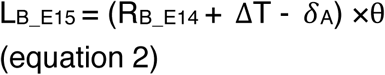

We solve for δA using the observed L_A_E15_ (equation 1), then use the constraint δ_A_ + _B_ = ΔT to obtain δB, and then use equation 2 to ***predict*** L_B_E15_. The predicted value is then compared to the independently observed L_B_E15_ (MMV, see table) as a test of the model.

Substituting all the known values (L_A_E15_, R_B__E14, ΔT, andθ; from MMV), we find δ_A_:

δ_A_= 0.127 mm

Using this we find δ_B_ = 0.108 mm

Therefore:

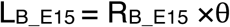

L_B_E15_ = 6.324 mm

This calculated value is within 2% of the observed value for L_B_E15_ obtained from MMV (see table). A reverse calculation to estimate L_A_E15_ also falls within 2% of the true observed value.

A case where both surfaces shift equally (δ_A_ = δ_B_ = ΔT/2 ≈ 0.118 mm each) was also calculated (not shown) where the error falls even further (to within 1%).

## Conclusion

Model 1 overpredicts both L_A_E15_ (by 14%) and L_B_E15_ (by 7%). Model 2, fit to L_A_E15_, predicts L_B_E15_ within 2%. The bidirectional model is therefore strongly supported, with apical and basal contributions approximately 54% and 46% of total thickness change respectively.

## Full-width-at-half-maximum (FWHM) Measurement and Mean corrected total cell fluorescence (CTCF) (Fig. 4B)

### FWHM

The measurements for FWHM to quantify the f-actin cell-cell junction thickness using phalloidin, a previously reported method [36] was employed.

1. Draw a line (using the straight-line tool in image J) on the cell-cell junction border. The line drawn was at least 2-3 times the width of the junction.
2. Draw the plot profile (Analyze > Plot profile or Ctrl+K). 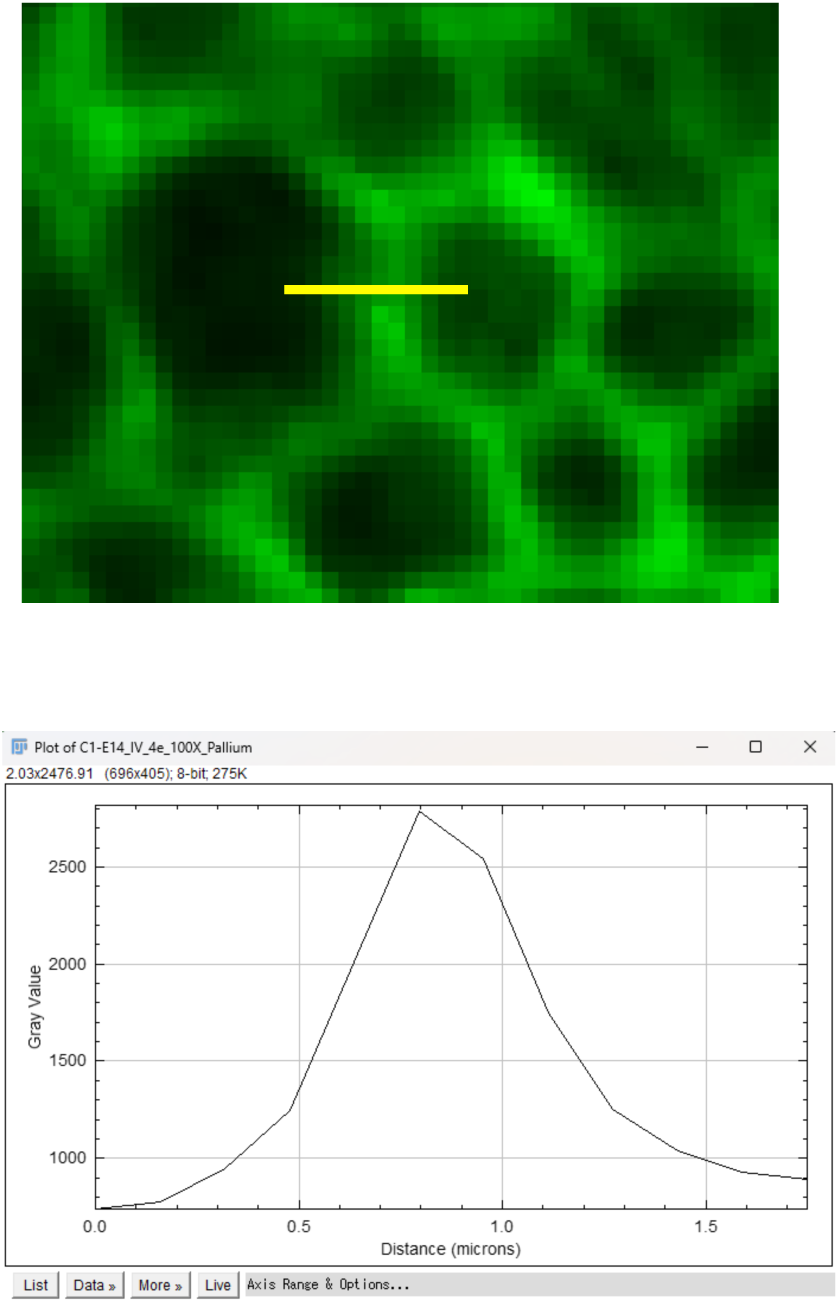
3. Click on the **list** button on the plot profile to get a list of values. Copy the list. 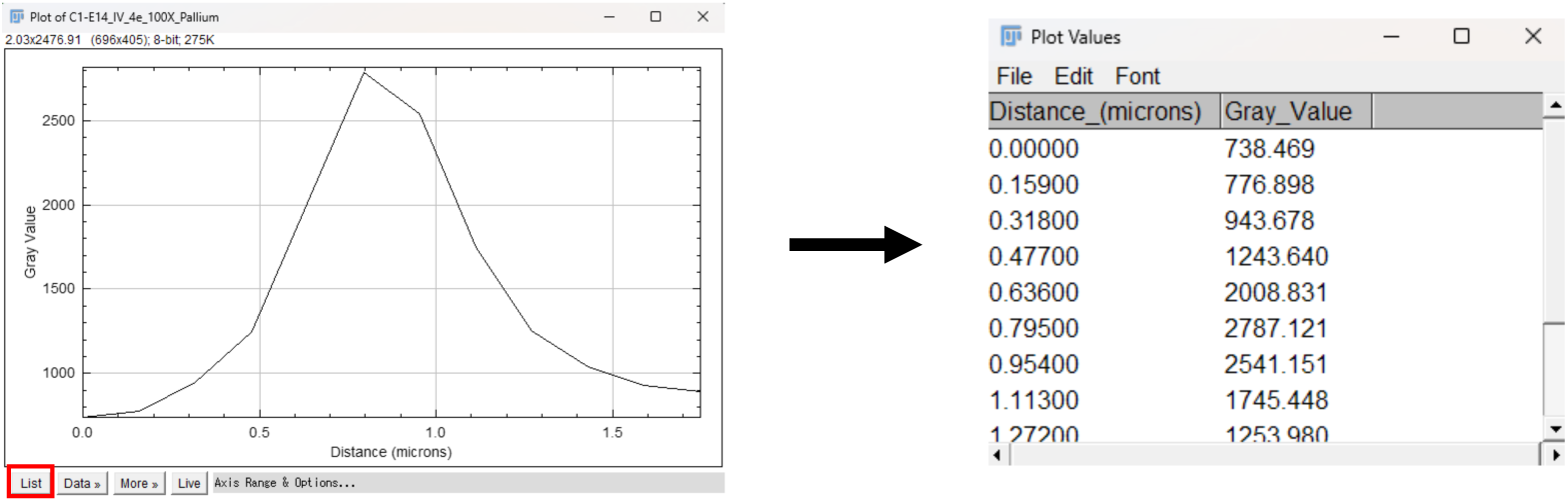
4. Search for ‘Curve fitting’ or go to (Analyze > Tools > Curve fitting). Paste the copied list here. In the drop-down menu select ‘Gaussian’ and click ‘Apply’. 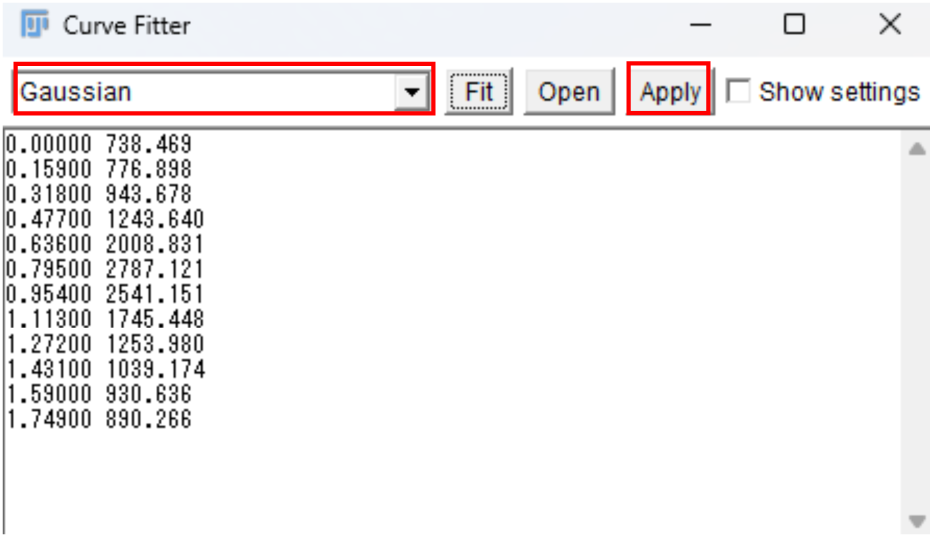
5. From the Gaussian fit 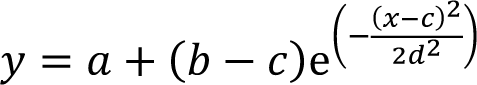 extract parameters a (baseline), b (peak amplitude), and d (width), then compute FWHM as: 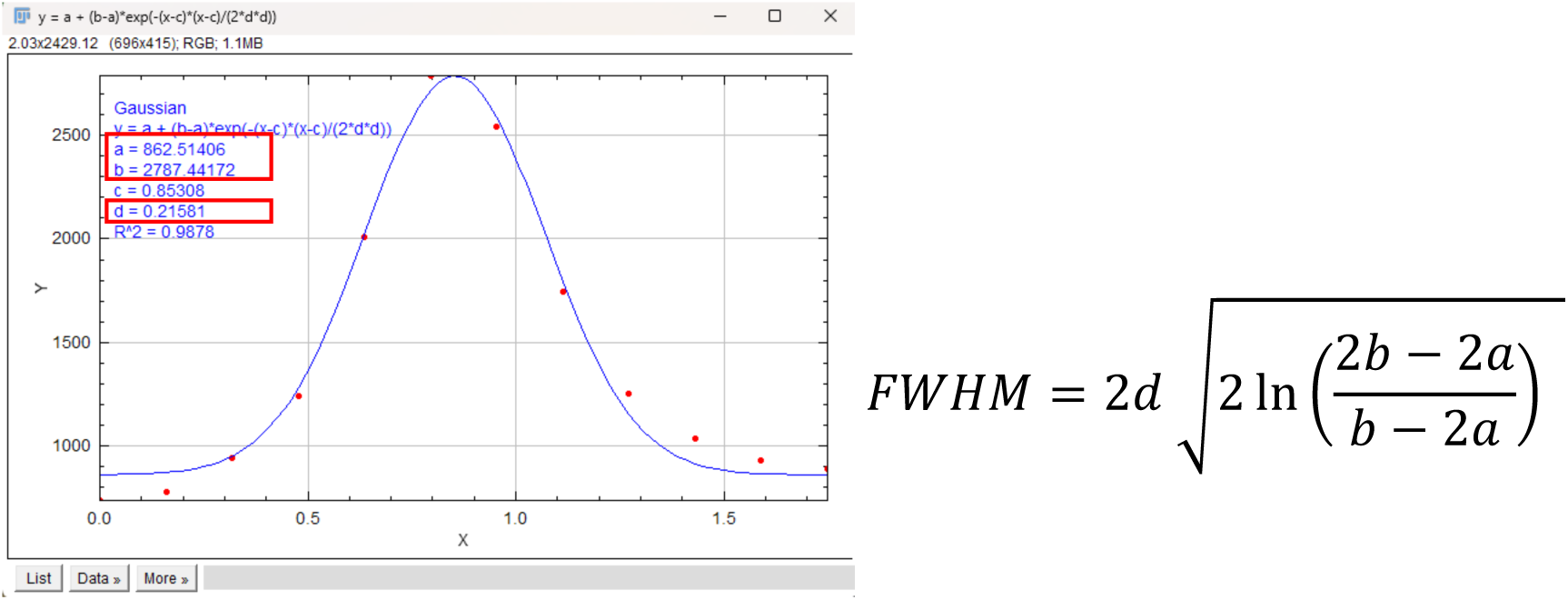

### Mean CTCF

For the quantification of pMLC in the apices of pallium and GE, mean corrected total cell fluorescence (Mean CTCF) was used. Unlike F-actin (visualized with phalloidin) or junction-associated proteins such as ZO-1, which are sharply enriched at cell-cell boundaries, pMLC distribution within individual cells was relatively uniform. Whole-cell pMLC levels reflect the total pool available for junctional contractility, and differences in whole-cell pMLC intensity between regions therefore serve as a proxy for differences in junctional contractility-generating capacity.

Co-staining with phalloidin or ZO-1 clearly demarcated the cell boundary (Fig., top), enabling unambiguous segmentation of individual cells. By contrast, pMLC signal was distributed throughout the cell body without sharp enrichment at the junction (Fig., bottom), making junction-restricted quantification unreliable. We therefore measured Mean CTCF over the whole cell body, defined by the phalloidin/ZO-1 boundary, allowing direct comparison of pMLC levels between pallium and GE without bias from junction-segmentation choices.

First, CTCF was calculated as: Integrated Density − (Cell Area × Mean Background Intensity) (in ImageJ/Fiji). Background values were obtained by averaging three representative cell-free areas in the image. To normalize for variations in cell size, the CTCF was divided by the area of the selected cell to determine the Mean CTCF. A total of 20-22 cells were analyzed for each embryo (*n = 4* embryos for pallium, *n = 4* for GE). The averaged Mean CTCF for each embryo was used in statistical analysis.

**Fig.**
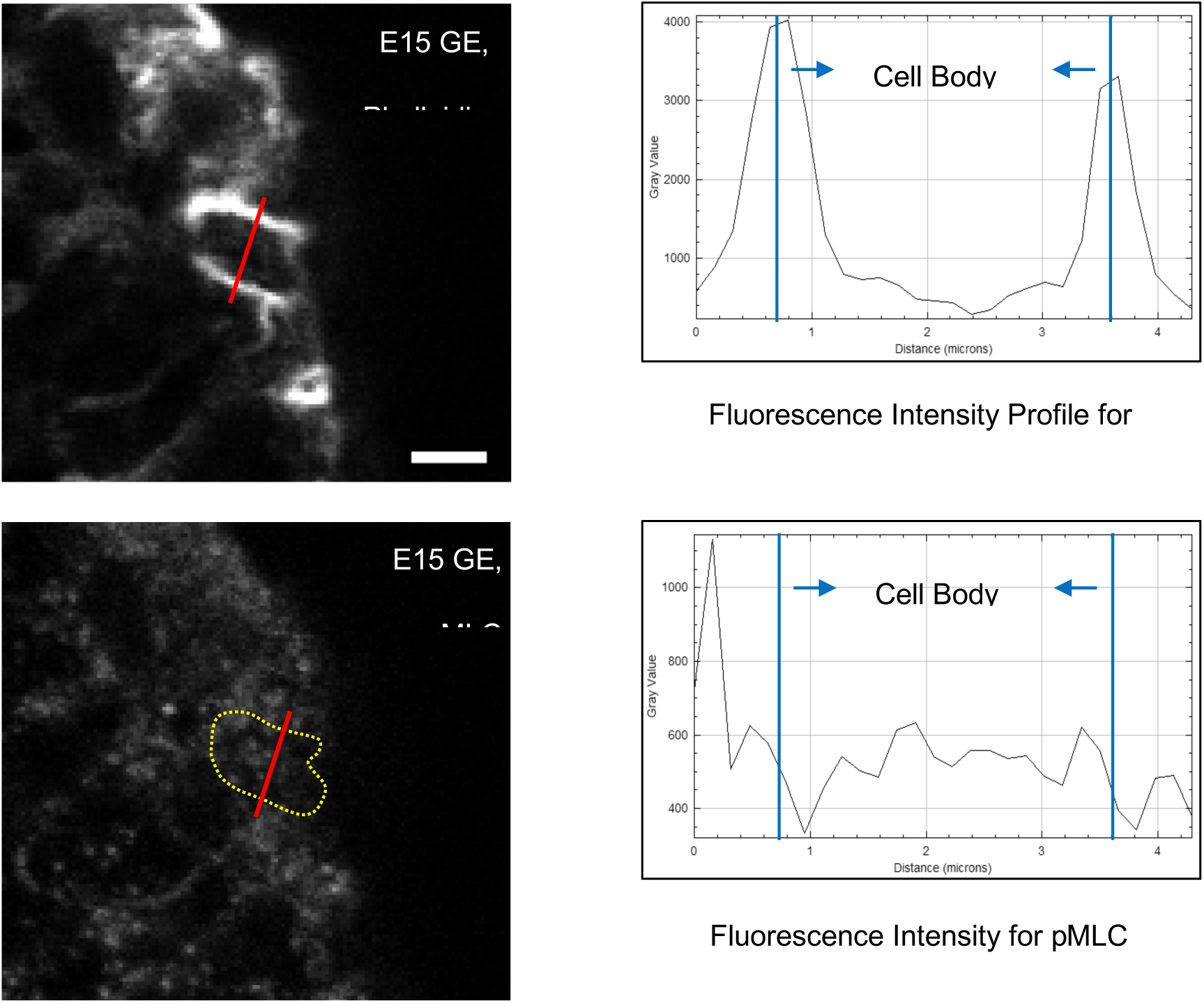
Whole-cell pMLC quantification. Top: Phalloidin staining (left) and corresponding intensity profile (right) along the indicated line ROI (red, left panel) shows two sharp peaks at the cell-cell junctions, clearly demarcating the cell boundary. Bottom: pMLC staining in the same cell (left; cell boundary outlined in yellow, inferred from phalloidin) and corresponding intensity profile (right) shows a relatively uniform distribution across the cell body, lacking sharp junctional peaks. Y-axes differ between panels to accommodate signal range. E15 GE; scale bar: 2 μm.

## Supplemental Movies

**Movie 1.**
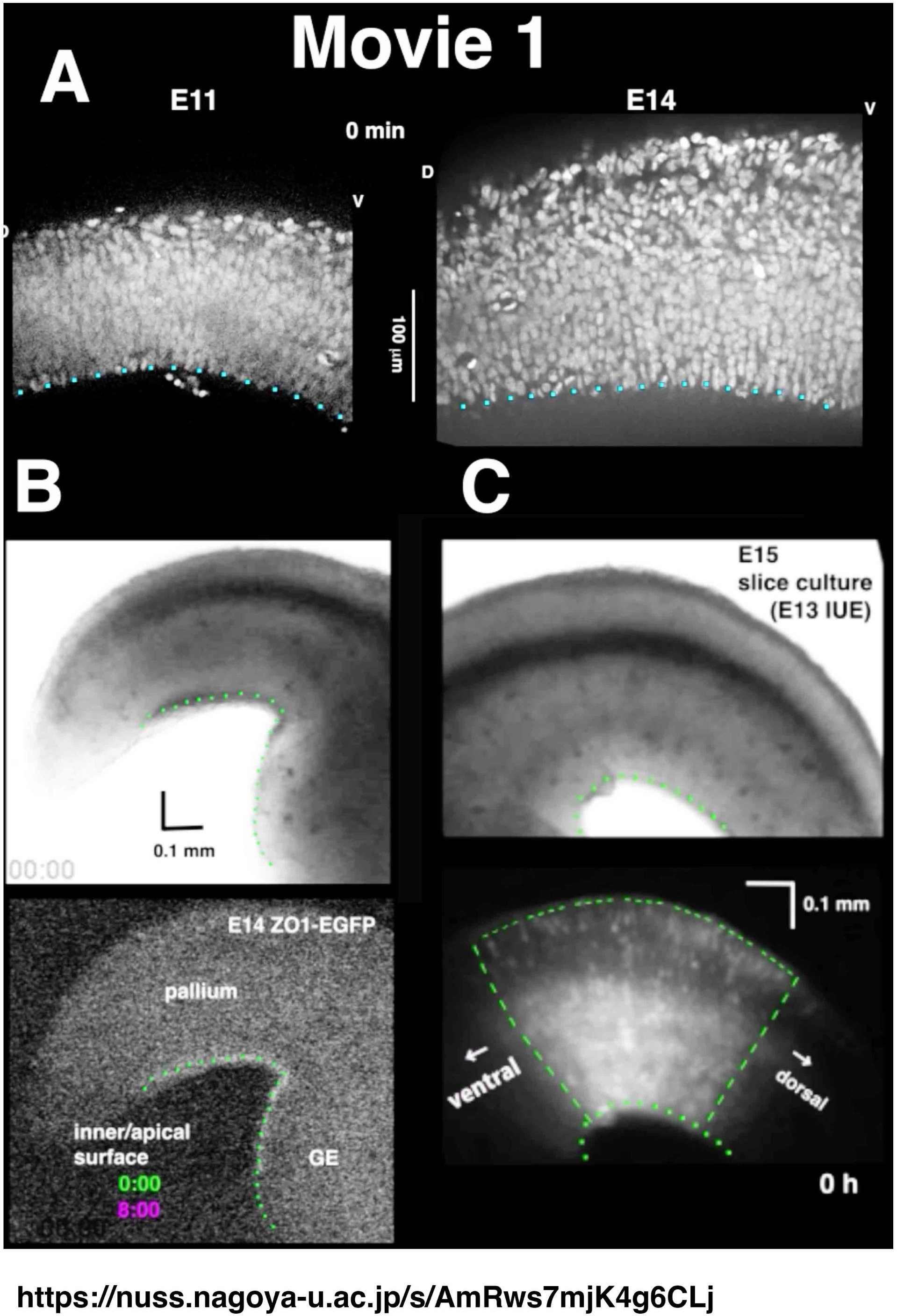
Inward thickening of cultured midembryonic cerebral pallial walls. A) Comparative time-lapse imaging of E11 and E14 pallial wall slices (coronally prepared from H2B-mCherry mice) for 6 h. D, dorsal. V, ventral. B) Bright-field and fluorescent time-lapse imaging of a cerebral wall slice prepared from an E14 ZO1-EGFP mouse. GE, ganglionic eminence. C) Bright-field and fluorescent time-lapse imaging of a cerebral wall slice prepared from an E15 ICR mouse that had been electroporated with an EGFP-expressing plasmid vector at E13.

**Movie 2.**
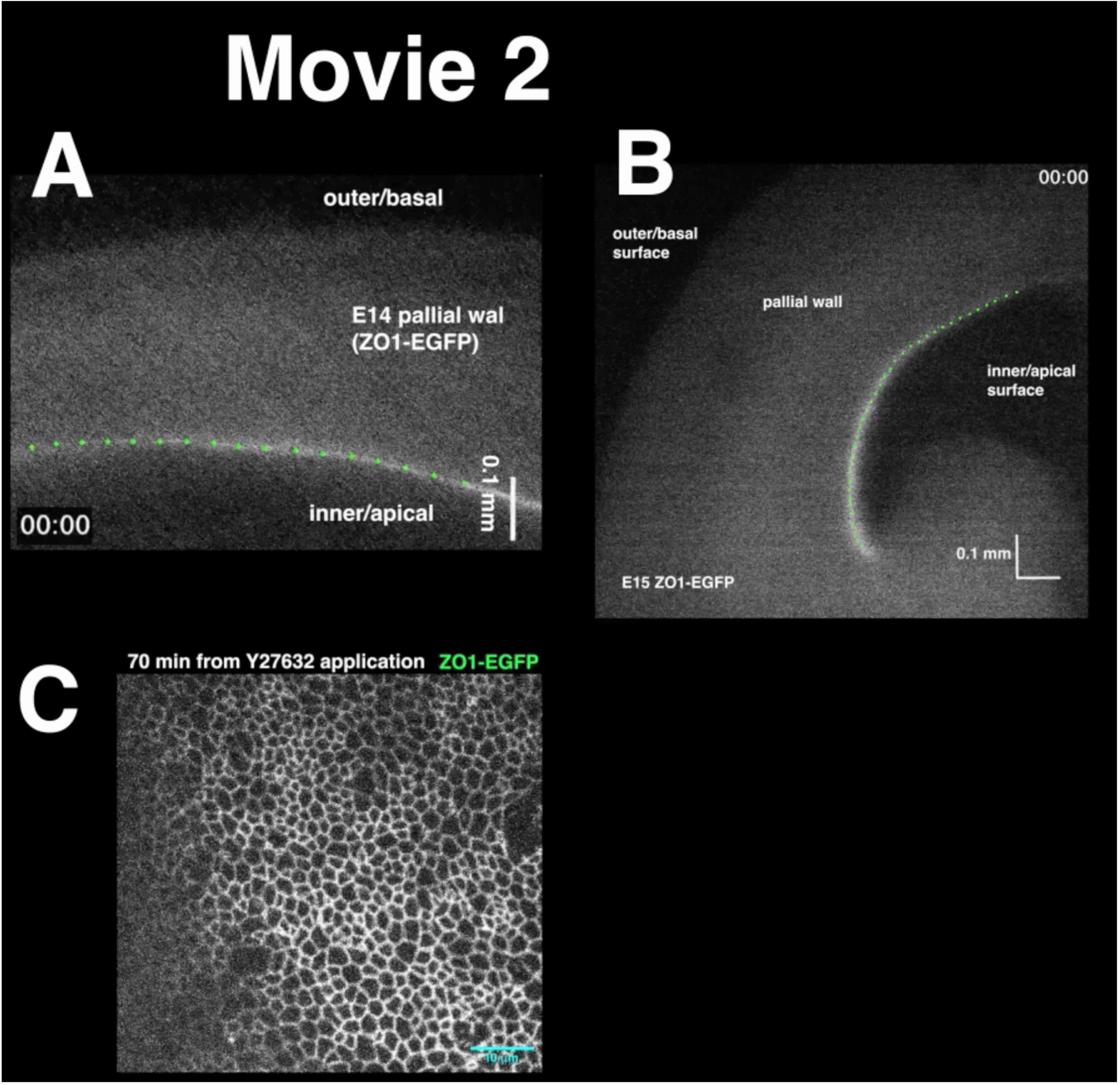
Responses of cultured midembryonic pallial walls to Y-27632. A) A pallial wall slice prepared from an E14 ZO1-EGFP mouse. Y-27632 addition caused the apical surface to cease the inward/apical-ward shift, followed by its slight retraction to the opposite side. B) A ZO1-EGFP-labeled pallial wall prepared at E15. Y-27632 application resulted in waviness of the apical surface. C) *En face* high-magnification imaging of a pallial wall prepared from an E14 ZO1-EGFP mouse. Big holes much larger than the size of normal meshes (endfeet) were formed.

**Movie 3.**
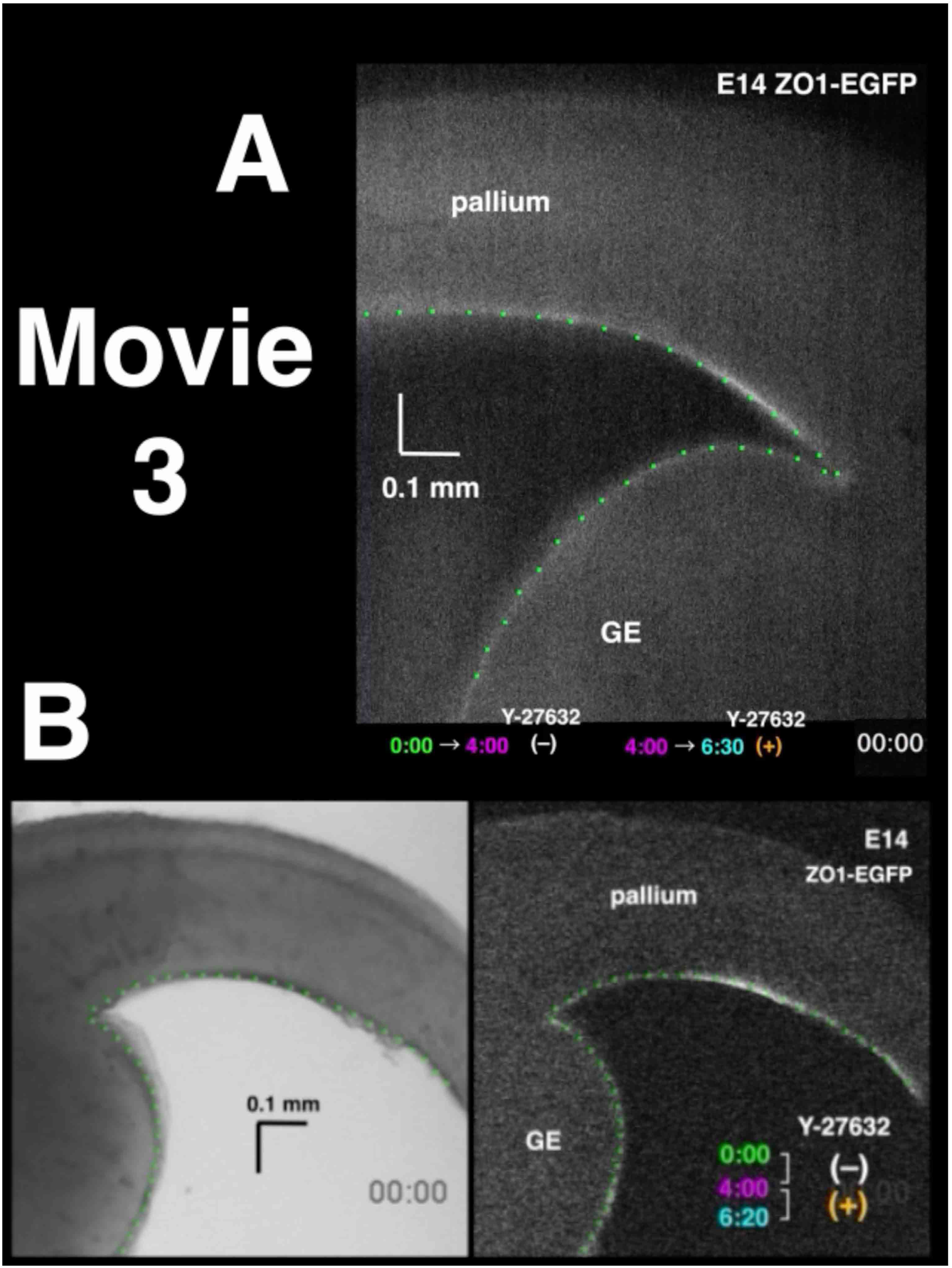
Responses of cultured midembryonic GE walls to Y-27632. A) A pallial wall and its ventral neighbor ganglionic eminence (GE) prepared from an E14 ZO1-EGFP mouse. While Y-27632 addition caused the pallial apical surface to cease the inward/apical-ward shift and retract to the opposite side, the GE apical surface continued to shift inward even after Y-27632 addition. B) Bright-field and fluorescent time-lapse imaging of a cerebral wall slice prepared from an E14 ZO1-EGFP mouse. Yellow arrowhead, slight bulging of GE in response to Y-27632 addition.

## Notes

### Competing Interest Statement

The authors have declared no competing interest.

